# Bacterial glycocalyx integrity drives multicellular swarm biofilm dynamism

**DOI:** 10.1101/2020.09.30.318626

**Authors:** Fares Saïdi, Nicolas Y. Jolivet, David J. Lemon, Arnaldo Nakamura, Anthony G. Garza, Frédéric J. Veyrier, Salim T. Islam

## Abstract

Bacterial surface exopolysaccharide (EPS) layers are key determinants of biofilm establishment and maintenance, leading to the formation of higher-order 3D structures conferring numerous survival benefits to a cell community. In addition to a specific EPS glycocalyx, we recently revealed that the social δ-proteobacterium *Myxococcus xanthus* secretes a novel biosurfactant polysaccharide (BPS), with both EPS and BPS polymers required for type IV pilus (T4P)-dependent swarm expansion via spatio-specific biofilm expression profiles. Thus the synergy between EPS and BPS secretion somehow modulates the multicellular lifecycle of *M. xanthus*. Herein, we demonstrate that BPS secretion functionally-activates the EPS glycocalyx via its destabilization, fundamentally altering the characteristics of the cell surface. This impacts motility behaviours at the single-cell level as well as the aggregative capacity of cells in groups via EPS fibril formation and T4P assembly. These changes modulate structuration of swarm biofilms via cell layering, likely contributing to the formation of internal swarm polysaccharide architecture. Together, these data reveal the manner by which the interplay between two secreted polymers induces single-cell changes that modulate swarm biofilm communities.

## INTRODUCTION

The detection of glycocalyces surrounding bacterial cells remains a seminal discovery in bacterial physiology, giving rise to the biofilm concept for surface-attached microbial community growth within a polysaccharide matrix (1). Within a biofilm, bacteria can physically interact, be protected from external stressors (e.g. antibiotics, reactive oxygen species, dehydration, etc.), replicate, communicate via secreted signals, and differentiate their functions (2, 3). While the importance of secreted polysaccharides for biofilm formation is widely appreciated, the mechanisms by which these polymers promote 3D matrix structuration and the cellular organization within are areas of intense study (4).

Robust biofilm existence is exemplified by the social multicellular lifecycle of *Myxococcus xanthus*, a predatory Gram-negative δ-proteobacterium (2, 5, 6). Groups of *M. xanthus* cells are encased within a secreted polysaccharide matrix, promoting intimate contacts. On surfaces, swarms of such cells are able to cooperatively predate prey microorganisms, saprophytically feeding on the degradation products. When nutrients become scarce, *M. xanthus* cells within a swarm biofilm secrete a signalling molecule that accumulates to a certain local threshold above which the developmental program is initiated, leading to the aggregation of thousands of cells and the formation of fruiting bodies. Functional differentiation within the swarm leads to three subpopulations, namely (i) myxospore-forming cells within the fruiting body lumen, (ii) peripheral rods that remain at the base of the fruiting body, and (iii) motile foragers that continue their outward trajectory from the initial aggregate (7).

Two motility systems are required to effectuate these complex physiological outcomes, with each being differentially active depending on the nature of the substratum. On hard surfaces, gliding (i.e. “adventurous” [A]) motility predominates, mediated by substratum coupling and directed transport of the trans-envelope Agl–Glt machinery at bacterial focal adhesion (bFA) sites (8-10). On soft substrata, cell groups move via type IV pilus (T4P)-dependent (i.e. “social” [S]) motility (11, 12). As a result of these complimentary systems, *M. xanthus* forms highly structured, yet dynamic, biofilms. On hard substrata, *M. xanthus* swarm biofilms expand along a radial vector as well as vertically away from the substratum, resulting in the formation of stratified cell layers (13, 14). Cells within each layer are motile, densely packed, and aligned along their long axes, displaying the properties of an active nematic liquid-crystal state of matter (15). This stratification on hard surfaces may require a functional gliding motility apparatus capable of coupling to an external contacting surface (e.g. the substratum and/or adjacent cells) as layer formation is severely compromised when the substratum-coupling adhesin of the Agl–Glt apparatus (CglB) is absent (10, 13). Prolonged incubation in such biofilms can lead to cells becoming connected via a network of outer-membrane vesicle (OMV) chains and OM tube (OMT) projections (16). In contrast, knowledge of internal swarm architecture on soft substrata is more limited. Of note, the leading edge of such swarms was shown via electron microscopy to contain discreet bundles of aligned cells, encased in Ruthenium Red-stained structures termed polysaccharide “microchannels” (17).

Several long-chain sugar polymers are synthesized by *M. xanthus* in order to modulate its complex lifecycle (18). For cells in development undergoing sporulation, the <u>ma</u>jor <u>s</u>pore <u>c</u>oat (MASC) polymer is produced to surround myxospores in a protective layer (19, 20). For non-sporulating cells, motility is affected by O-antigen-capped LPS (21-23), as well as a poorly understood “slime” polymer that is proposed to promote adhesion of the Agl–Glt gliding complex to the underlying surface, and which is left behind in trails following transit of gliding cells (24, 25). In addition, the bacterium synthesizes <u>e</u>xo<u>p</u>oly<u>s</u>accharide (EPS); this is a specific secreted sugar polymer required for T4P-dependent swarm spreading, which inhibits natural transformation and constitutes the principal matrix polysaccharide in *M. xanthus* biofilms (26-31). Recently, we reported that *M. xanthus* also synthesizes and secretes a novel <u>b</u>iosurfactant <u>p</u>oly<u>s</u>accharide to the extracellular milieu that is essential for T4P-dependent swarm spreading (32). Within an expanding swarm biofilm, BPS biosynthetic machinery is more highly expressed in the swarm centre, whereas EPS biosynthetic machinery is more highly expressed at the swarm periphery, pointing to spatially-distinct roles for each polysaccharide in the maturation of multicellular swarm biofilms (32).

Each of EPS, BPS, and MASC is synthesized by a separate Wzx/Wzy-dependent pathway, with each respective component given the suffix X (e<u>x</u>opolysaccharide), B (<u>b</u>iosurfactant), or S (<u>s</u>pore coat) (32-35). Individual polysaccharide repeat units in such pathways are assembled on the lipid carrier undecaprenyl pyrophosphate (UndPP) at the cytoplasmic leaflet of the inner membrane (IM), followed by processing via a suite of integral IM proteins (36). Repeat units bound to UndPP are first transported across the IM by the Wzx flippase (37-40). UndPP-linked repeats in the periplasmic leaflet of the IM are then polymerized by Wzy (41-43), to modal lengths specified by Wzz/Wzc polysaccharide co-polymerase (PCP) proteins (41). BPS-pathway WzcB is of the PCP-2A class (44) as it contains an attached cytosolic bacterial tyrosine autokinase (BYK) domain (32, 45, 46). Conversely, EPS- and MASC-pathway WzcX and WzcS (respectively) are of the PCP-2B class as they do not encode a fused BYK domain; instead, these pathways encode standalone WzeX/S BYK proteins (32, 47) for association with their cognate WzcX/S PCP. In turn, Wzb bacterial tyrosine phosphatase (BYP) proteins are also encoded to control the phosphorylation state of PCP-2A Wzc proteins and PCP-2B-associated Wze proteins (48). The *M. xanthus* Wzb BYP (PhpA) has been shown to dephosphorylate BYK WzeS as well as the BYK domain of WzcB (49), implicating Wzb in MASC and BPS biosynthesis. The cytosolic (de)phosphorylation states of PCP-2A Wzc and PCP-2B-associated Wze proteins control not only polymer modal length modulation, but also secretion of the respective heteropolysaccharide across the OM via the Wza translocon (50, 51).

A range of activators and inhibitors are known to impact *M. xanthus* EPS biosynthesis (reviewed in (18)), with the Dif chemosensory pathway the most noteworthy (52-54). Positive regulation of EPS production is mediated by (i) the methyl-accepting chemotaxis protein DifA, (ii) the CheW-like coupling protein DifC, (iii) the CheA-like histidine kinase DifE, and (iv) the EpsW response regulator (phosphorylated by DifE) (55-57) Conversely, EPS production is negatively regulated by (i) the DifD response regulator, (ii) the CheC-like phosphatase DifG, and (iii) Nla19, an NtrC-like transcriptional regulator (54, 58). Despite these details, the specific functional links between the Dif pathway and the Wzx/Wzy-dependent EPS biosynthesis system have yet to be identified.

While the biosurfactant nature of BPS was previously demonstrated, the mechanism by which it promotes swarm structuration and T4P-dependent spreading was not known (32). Herein, we provide evidence/demonstrate that unlike conventional biosurfactants — which typically function as wetting agents secreted at colony fronts to condition the substratum and promote spreading — a major role of BPS is to change the activation state of the EPS surface glycocalyx. This impacts cell-level and community-scale behaviours leading to altered multicellular outcomes.

## RESULTS

### *BPS-deficient* M. xanthus *cells are hyper-aggregative*

Though BPS^−^ and EPS^−^ swarms were previously shown to be compromised for T4P-dependent swarm expansion — with the former elaborating a fuzzy morphology and the latter a smooth phenotype — under nutrient limitation BPS^−^ cells were still able to aggregate and form spore-filled fruiting bodies, similar to WT (but not EPS^−^) swarms (32) (**Fig. 1**). However, the developmental transition to fruiting bodies for BPS^−^ swarms could take place at lower initial cell densities compared to WT swarms, suggesting that cells may aggregate more efficiently in BPS^−^ swarms (32). To specifically probe aggregative differences among *M. xanthus* cell-surface polysaccharide mutants, the real-time ability of cells to auto-aggregate in liquid media was compared, a phenomenon known to be dependent on the presence of cell-surface EPS and an extendable T4P (22, 26, 30, 33, 55, 56, 58-68). While both EPS^−^ and T4P^−^ cells remained in suspension, BPS^−^ cells rapidly auto-aggregated by the first post-mixing time point (10 min), resulting in faster initial unaided sedimentation relative to WT cells; after ∼40 min, WT and BPS^−^ cells in rich media continued to sediment, but at comparable rates, consistent with continued metabolism (**Fig. 2A**). The same rapid auto-aggregation phenotype at the same post-mixing time point (10 min) was observed for BPS^−^ cells (relative to WT) under non-metabolizing conditions in minimal buffer, with the auto-aggregation levelling off at ∼40 min (**Fig. 2B**). Together, these cell-level hyper-aggregation data (i) help explain fruiting body formation for BPS^−^ swarms at lower cell densities (32), (ii) suggest that T4P may indeed be extendable in BPS^−^ cells, and (iii) are consistent with differences in the general surface properties of BPS^−^ cells.

**Figure 1.**
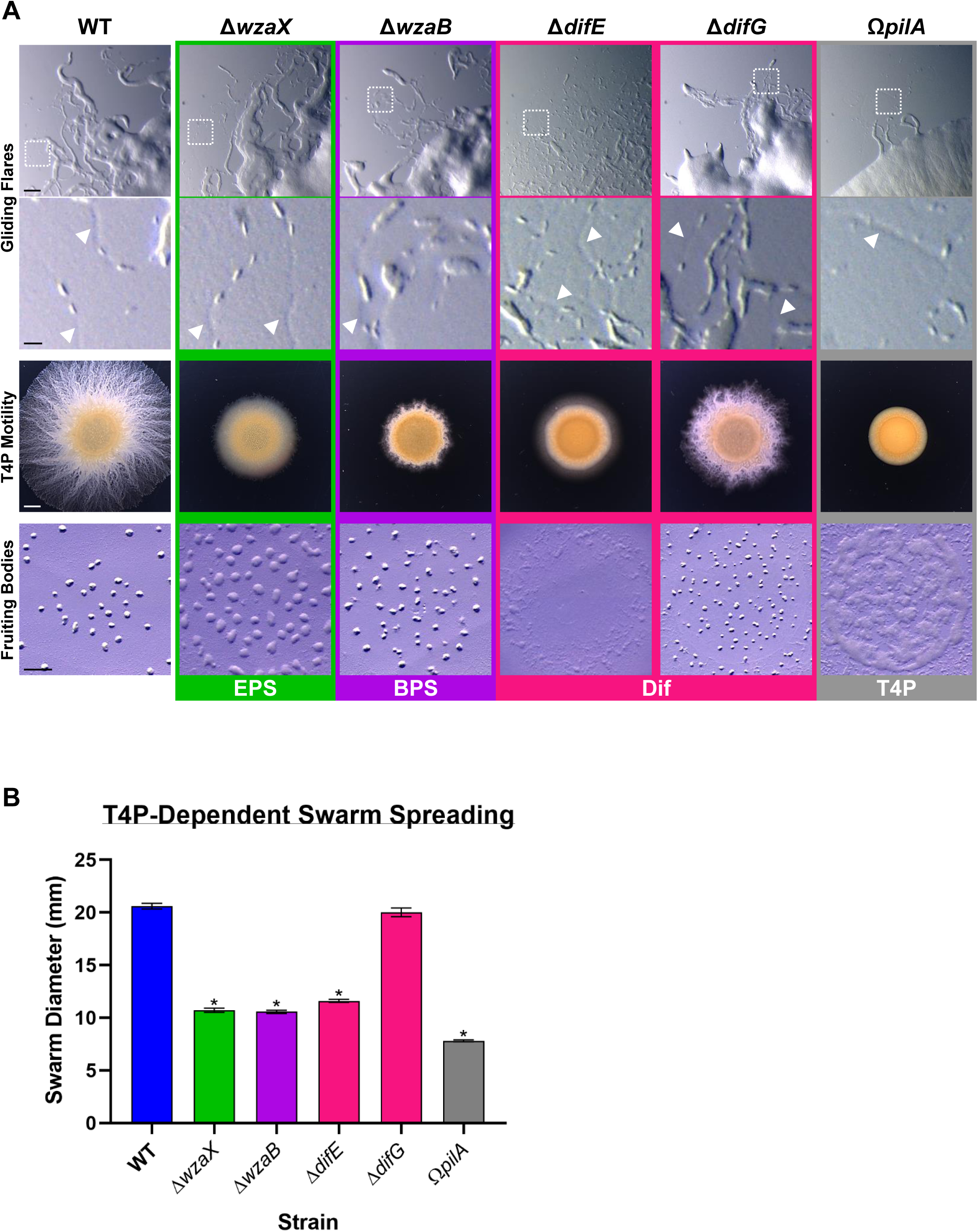
**(A)** Motility and developmental phenotypes for various mutants with altered levels of secreted polysaccharides. Top panels (upper): gliding motility flares on CYE 1.5% agar after 30 h (scale bar: 50 µm). Top panels (lower): magnified view of white hatched box in corresponding upper panel showing furrows in the agar substratum containing transiting cell groups (scale bar: 10 µm). Arrowheads (⊳) indicate furrows in the agar left by previously-transited cells and/or cell groups, revealed by extreme oblique illumination of agar surface. Middle panels: T4P-dependent swarm spreading on CYE 0.5% agar after 72 h (scale bar: 2 mm). Bottom panels: fruiting body formation on CF 1.5% agar after 75 h (scale bar: 1 mm). **(B)** Bar graphs of diameters of swarms grown on CYE 0.5% agar for T4P-dependent motility at 72 h. For each strain, the mean value of 3 biological replicates (+/− SEM) is plotted. Asterisks (*) denote datasets displaying statistically significant differences (*p* < 0.0001) relative to both WT and Δ*difG* strains, as determined via unpaired two-tailed Student’s t-test analyses.

**Figure 2.**
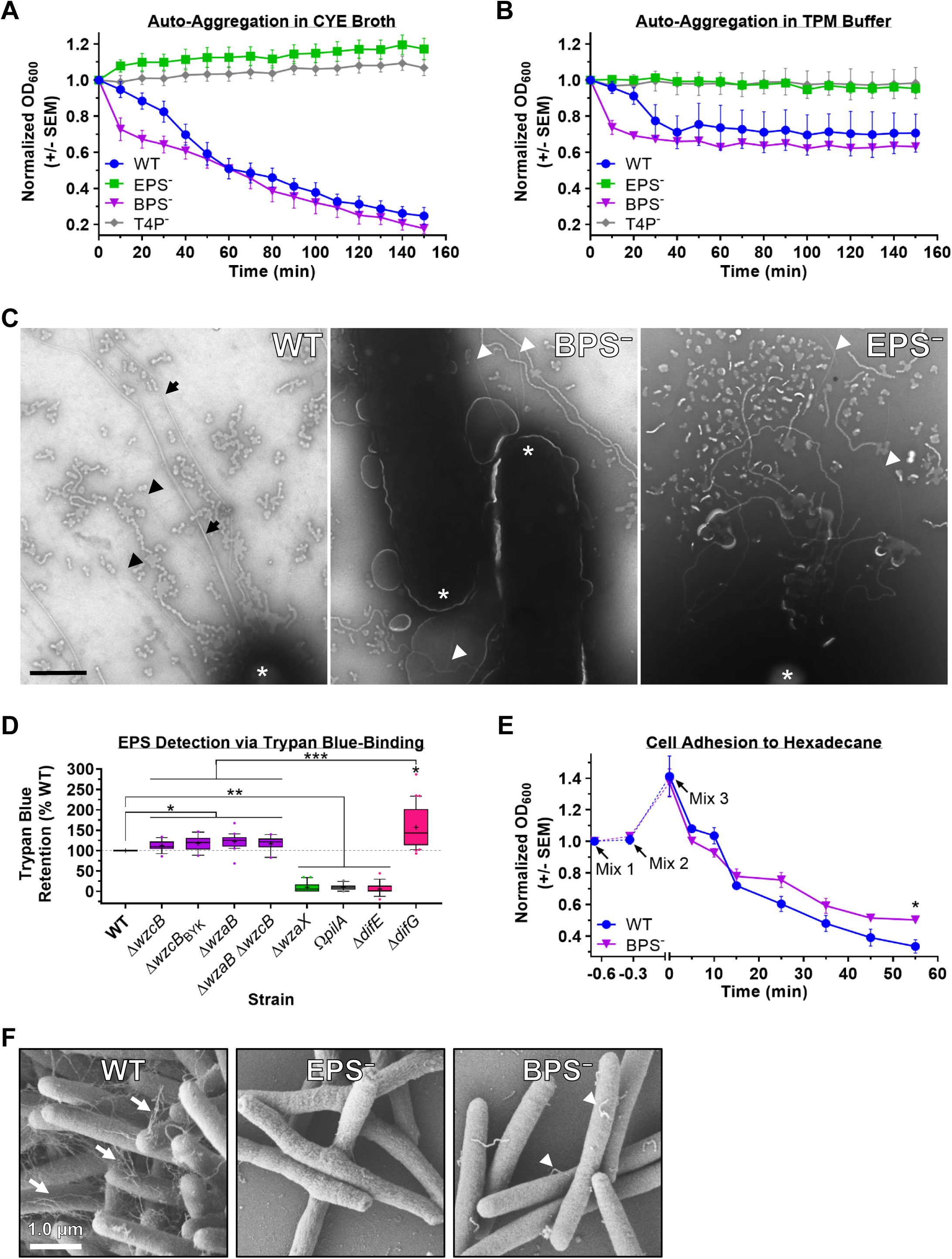
Auto-aggregation profiles of WT, EPS^−^ (Δ*wzaX*), BPS^−^ (Δ*wzaB*), and Ω*pilA* strains resuspended in **(A)** CYE rich medium (mean values of 6, 6, 6, and 5 biological replicates [+/− SEM], respectively) and **(B)** TPM minimal buffer (mean values of 3 biological replicates [+/− SEM]). **(C)** Representative transmission electron micrographs of WT, BPS^−^ (Δ*wzaB*), and EPS^−^ (Δ*wzaX*) cells on copper grids, taken at 10 000× magnification (scale bar: 500 nm). Arrowheads (⊳) denote thin (likely single) T4P filaments. Arrows (⇒) denote thick (likely bundled) T4P filaments. Asterisks (*) denote cell poles. **(D)** Boxplots of Trypan Blue dye retention to indicate the levels of EPS production in various strains relative to WT. The lower and upper boundaries of the boxes correspond to the 25^th^ and 75^th^ percentiles, respectively. The median (line through centre of boxplot) and mean (+) of each dataset are indicated. Lower and upper whiskers represent the 10^th^ and 90^th^ percentiles, respectively; data points above and below the whiskers are drawn as individual points. Asterisks denote datasets displaying statistically significant differences in distributions (*p* < 0.05) shifted higher (**) or lower (*) than WT, as determined via Wilcoxon signed-rank test performed relative to “100”. Data for Δ*difE* and Δ*difG* was heretofore unreported and acquired at the same time as the published values for other strains (32), reproduced with permission. **(E)** Microbial adhesion to hydrocarbons (MATH) test of WT and BPS^−^ (Δ*wzaB*) strain binding to hexadecane; values are the mean of 3 biological replicates (+/− SEM). Mix 1, initial pellet resuspension in CYE; Mix 2, supplemental control mixing; Mix 3, mixing upon addition of hexadecane (*t* = 0). Asterisk (*) denotes dataset displaying statistically significant difference in time point mean value (*p* = 0.0176) compared to WT, as determined via unpaired Student’s t test. **(F)** Scanning electron micrographs of WT, EPS^−^ (Δ*wzaX*), and BPS^−^ (Δ*wzaB*) cells, taken at 20 000× magnification (scale bar: 1.0 µm). Arrows denote EPS fibrils connecting cells. Arrowheads denote potential surface-associated fibril material not engaged in inter-cell connections.

### *BPS-deficient* M. xanthus *cells can extend T4P*

To directly probe T4P assembly, WT, BPS^−^ and EPS^−^ cells from liquid culture were analyzed via transmission electron microscopy (TEM). As expected, from the poles of WT cells, numerous T4P projections could be observed, including both thin filaments (likely single pili) and thick filaments (likely bundles of pili); the presence of T4P bundles is supported by the observation that at certain points along the thick filament, branching occurred, with the new offshoots resembling thin filaments (**Fig. 2C**). While no EPS^−^ cells were observed with attached T4P projections emanating from a cell pole, thin T4P-like filaments were detected on the grid (**Fig. 2C**). This may suggest that EPS^−^ cells can still assemble a T4P, but that the presence of cell-surface EPS contributes to strengthening the apparatus. Intriguingly, BPS^−^ cells were observed to extend T4P, but they were not as prevalent as in WT cells; moreover, the T4P filaments detected in BPS^−^ cells were typically shorter than those in WT cells, and not found in presumed thick T4P bundles (**Fig. 2C**). Thus while BPS^−^ cells can still extrude T4P projections, the assembly of these apparatus appears to be compromised relative to WT cells, helping to explain the deficiency in T4P-dependent motility in BPS^−^ cells (**Fig. 1**) (32).

### Increased Trypan Blue-retention by BPS^−^ cells is inconsistent with EPS overproduction

Retention of Trypan Blue dye is used as a readout for *M. xanthus* cell-surface EPS levels (30, 58, 69). Intriguingly, despite compromised T4P-dependent swarm spreading (32, 33), BPS-pathway mutants Δ*wzcB*, Δ*wzcB*_BYK_, Δ*wzaB*, and Δ*wzaB* Δ*wzcB* — in which periplasmic BPS polymerization should be permitted but secretion compromised — reproducibly bound more Trypan Blue than WT cells (combined interquartile range of 102–133% of WT) (**Fig. 2D**) (32). This dye-retention difference was manifested despite equivalent levels of the same cell-associated EPS sugars detected in WT and BPS^−^ cells (32). However, the abovementioned four BPS-pathway mutant strains still bound significantly less Trypan Blue than the Δ*difG* strain (interquartile range of 113–202% of WT) (**Fig. 2D**). The latter is a mutant in the Dif chemosensory pathway in which EPS production is no longer negatively-regulated, resulting in increased EPS production; this is compared to a Δ*difE* strain in which EPS production is downregulated (**Fig. 2D**) (55, 56, 58, 70). In fact, EPS overproduction does not significantly compromise T4P-dependent swarm spreading relative to a BPS deficiency (**Fig. 1B**), which severely impairs swarm spreading (32). Taken together with the comparative dye-retention analyses of EPS regulatory mutants (**Fig. 2D**), these data are consistent with the elevated retention of Trypan Blue by the abovementioned BPS-pathway mutants not being due to EPS overproduction. Instead, this may point to differences in surface properties between WT and BPS^−^ cells.

### *BPS deficiency decreases the relative cell-surface hydrophobicity of* M. xanthus

To probe the physical properties of WT and BPS^−^ cell surfaces, both strains were subjected to MATH (microbial adhesion to hydrocarbons) testing to probe relative differences in cell-surface hydrophobicity (71). Cells of either strain were resuspended in (aqueous) liquid medium and mixed with the hydrocarbon hexadecane, after which the emulsion was allowed to clear (72). The rationale herein is that the greater the cell-surface hydrophobicity of a particular strain, the more cells removed from suspension through hydrophobic contacts with hexadecane, thus decreasing the turbidity of the suspension (71). Compared to OD_600_ readings taken before hexadecane addition-and-mixing, significantly fewer WT cells remained in suspension (i.e. lower OD_600_) following emulsion separation compared to BPS^−^ cells (**Fig. 2E**). These findings do not indicate that WT cells are hydrophobic per se, but rather that the WT cell surface is relatively more hydrophobic compared to that of BPS^−^ cells. Therefore, BPS secretion increases the relative hydrophobicity of the *M. xanthus* cell surface. Moreover, this dataset further supports the designation of BPS as a biosurfactant since biosurfactants have been extensively reported to change the relative cell-surface hydrophobicity of various bacterial cells (73). Finally, the reduced relative cell-surface hydrophobicity of BPS^−^ cells may help explain the higher-than-WT amounts of hydrophilic Trypan Blue binding observed for the Δ*wzcB*, Δ*wzcB*_BYK_, Δ*wzaB*, and Δ*wzaB* Δ*wzcB* mutants previously described (**Fig. 2D**) (32).

### BPS secretion is required for EPS surface-fibril formation

Given the multiple datasets pointing towards fundamental cell-surface differences between WT and BPS^−^ cells (**Fig. 2A-E**), surface morphologies for these strains (as well as EPS^−^ cells) were directly visualized via scanning electron microscopy (SEM). Consistent with previous reports, WT cells were shown to be connected via networks of EPS fibrils, while EPS^−^ cells lacked any such connections (16, 55, 63, 74-79) (**Fig. 2F**). Interestingly, while BPS^−^ cells still produce EPS (32) (**Fig. 2D**), no inter-cell fibril networks were observed for this strain (**Fig. 2F**). This may indicate that while EPS by itself is tightly held by individual cells, the secretion of BPS serves to sufficiently destabilize or loosen the cell-surface EPS glycocalyx, thus promoting fibril formation and inter-cell connections.

### EPS glycocalyx destabilization impacts single-cell behaviour

Since BPS^−^ cells are capable of forming fruiting bodies (**Fig. 1A**) (32), this suggests that BPS^−^ cells can still perform single-cell gliding motility, which is required for efficient fruiting body formation (80). For swarms grown on hard 1.5% agar, flare projections were observed emanating from the edge of the inoculated spot, a tell-tale sign of gliding motility by BPS^−^ cells (**Fig. 1A**). By way of severe oblique illumination of the samples, we were also able to visualize the furrow network left behind in the agar by lead gliding cells at the swarm edge (**Fig. 1A**). Previously revealed in detail by others using 3D profilometry, these physical depressions in the agar substratum were revealed to be the source of phase-bright trails classically attributed to slime deposition by *M. xanthus* cells gliding on agar (81). All strains tested produced furrows, indicating that the presence or absence of EPS and/or BPS does not qualitatively impact the formation of these substratum depressions. Moreover (while not possible to distinguish between single cells and cell groups), additional cells were detected following the path of the various furrows (**Fig. 1A**), supporting the notion of sematectonic stigmergic coordination for the phenomenon of trail following by *M. xanthus* cells on agar (82). At the single-cell level, the presence of a compacted surface glycocalyx in BPS^−^ cells, or the complete absence of this layer in EPS^−^ cells, resulted in faster gliding motility than in WT cells (**Fig. 3A**). Compared to BPS^−^ or EPS^−^ cells, this may indicate that the bulk volume of the destabilized EPS surface layer in WT cells adversely affects gliding efficiency.

**Figure 3.**
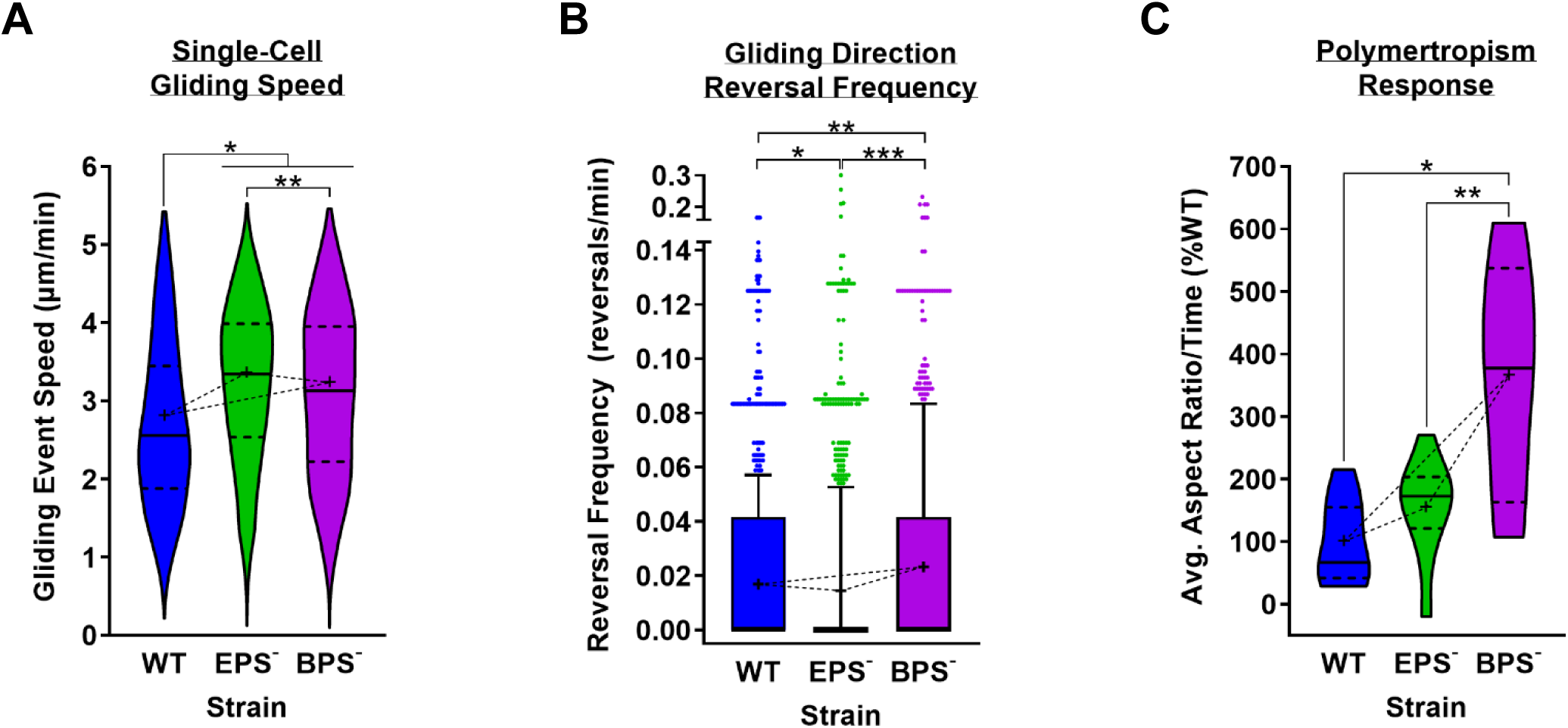
**(A)** Violin plots of single-cell gliding event speeds for WT, EPS^−^ (Δ*wzaX*), and BPS^−^ (Δ*wzaB*) cells on 1.5% agar pads (n = 2298 events across 4 biological replicates). A gliding event was defined as an instance of continuous translocation in a given direction. Cessation of motion by a given cell, followed by either a resumption of gliding in the same direction or a reversal of gliding direction, was considered the beginning of a new gliding event. The lower and upper boundaries of the plots correspond to the minimum and maximum values of the dataset, with the 25^th^ and 75^th^ percentiles displayed (*thick hatched black lines*). The median (*solid black line*) and mean (+) of each dataset are indicated. Asterisks denote datasets displaying statistically significant differences in distributions (*p* < 0.0001) between (*) WT and EPS^−^/BPS^−^ cells, as well as between (**) EPS^−^ and BPS^−^ cells, as determined via unpaired two-tailed Mann-Whitney test. **(B)** Boxplots of reversals per minute for tracked WT, EPS^−^, and BPS^−^ single cells on 1.5% agar pads (n = 1135 cells across 4 biological replicates). The lower and upper boundaries of the boxes correspond to the 25^th^ and 75^th^ percentiles, respectively. The median (line through centre of boxplot) and mean (+) of each dataset are indicated. Lower and upper whiskers represent the 10^th^ and 90^th^ percentiles, respectively; data points above and below the whiskers are drawn as individual points. Asterisks denote datasets displaying statistically significant differences in distributions between (*) WT and EPS^−^ cells (*p* = 0.0424), (**) WT and BPS^−^ cells (p < 0.0001), and (***) EPS^−^ and BPS^−^ cells (p < 0.0001), as determined via unpaired two-tailed Mann-Whitney tests. **(C)** Violin plots of polymertropism responses for WT, EPS^−^ (Δ*wzaX*), and BPS^−^ (Δ*wzaB*) polysaccharide secretion mutant strains. The lower and upper boundaries of the plots correspond to the minimum and maximum values of the dataset, with the 25^th^ and 75^th^ percentiles displayed (*thick hatched black lines*). The median (*solid black line*) and mean (+) of each dataset are indicated. Asterisks denote datasets displaying statistically significant differences in distributions between (*) WT and BPS^−^ swarms (*p* = 0.0006) and (**) EPS^−^ and BPS^−^ swarms (p = 0.0232), whereas distributions between WT and EPS^−^ swarms were not significantly different (p = 0.0845), as determined via unpaired two-tailed Mann-Whitney test. The number of biological replicates (*n*) used to analyze each strain as follows: WT (11), Δ*wzaX* (10), Δ*wzaB* (10).

We next probed the frequency at which single cells reversed their gliding direction. Cells were imaged at 30 s intervals for 50 frames. To avoid unintentionally lowering reversal frequency averages (by including cells tracked for a short time in which a reversal may not have yet manifested), we only analyzed cells continuously tracked for 30 or more frames. Cells deficient in EPS secretion were observed to reverse their gliding direction less frequently compared to WT cells (**Fig. 3B**), consistent with previous reports of lower reversal frequencies in Dif-pathway mutants in which EPS production is compromised (68, 83). Conversely, BPS^−^ cells were found to reverse their gliding direction more frequently than WT cells (**Fig. 3B**). Together, these data point to not only the general importance of EPS in regulating reversal frequency, but also its “activation state” as determined by the effects of BPS.

Given the differences in single-cell gliding behaviours described above, we examined the polymertropism responses of EPS^−^ and BPS^−^ cells. Polymertropism is a gliding motility-dependent process. It is measured via changes of the swarm aspect ratio, i.e. comparisons of changes in “east–west” expansion vs “north–south” expansion on an agar plate in response to the insertion of a small length of tubing between the edge of the agar and the “northern” wall of the Petri dish. The net effect of this agar compression is to align the polymers in the substratum matrix, allowing *M. xanthus* and other bacteria to preferentially spread in the “east–west” direction of the aligned substratum polymers (10, 84-86). While no significant differences in polymertropism responses were detected between WT and EPS^−^ swarms, BPS^−^ swarms demonstrated a remarkably enhanced capacity to spread in the “east–west” direction on compressed agar (**Fig. 3C**). This is the first known description of a hyper-polymertropic *M. xanthus* strain. While specific cellular factors contributing to the polymertropism response remain poorly understood, our data suggest that the presence of EPS as well as increased gliding speed may contribute to this enhanced “east–west” swarm expansion in response to mechanical changes in the substratum. The secretion of BPS thus affects *M. xanthus* behaviours at multiple levels of biological organization, from entire communities down to single cells.

### BPS secretion is required for cell stratification within swarms

We next sought to probe potential ultrastructural differences in swarm architecture leading to compromised T4P-dependent colony expansion. To highlight internal structures as per a previous report (17), spreading swarms were treated with Ruthenium Red, a polycationic dye that interacts with a range of polyanionic targets (87). This was carried out for swarms of WT, as well as the isogenic EPS^−^, BPS^−^, and MASC^−^ mutant strains, followed by negative-stain TEM of transversely-cut sections near the swarm edge (**Fig. 4, *inset***). This resulted in electron-dense labelling of the *M. xanthus* cell surface in the absence of EPS, BPS, or MASC secretion (**Fig. 4**). These analyses also revealed pronounced horizontal electron-dense structures separating stratified layers of WT and MASC^−^ cells, with such structures largely absent in BPS^−^ swarms and nonexistent in EPS^−^ swarms (**Fig. 4**). Vertical striations of this electron-dense material — connecting horizontal electron-dense structures above and below to form a self-contained so-called “microchannel” — were not detected (**Fig. 4**). The rod-shaped cells within the layered WT and MASC^−^ swarms, as well as the more irregularly-packed BPS^−^ swarm, were highly aligned along their long axes in the direction of migration, resulting in the round appearance of cells in the cross sections (**Fig. 4**). Cells in BPS^−^ swarms were also more closely-packed together compared to either WT, MASC^−^, or EPS^−^ swarms (**Fig. 4**).

**Figure 4.**
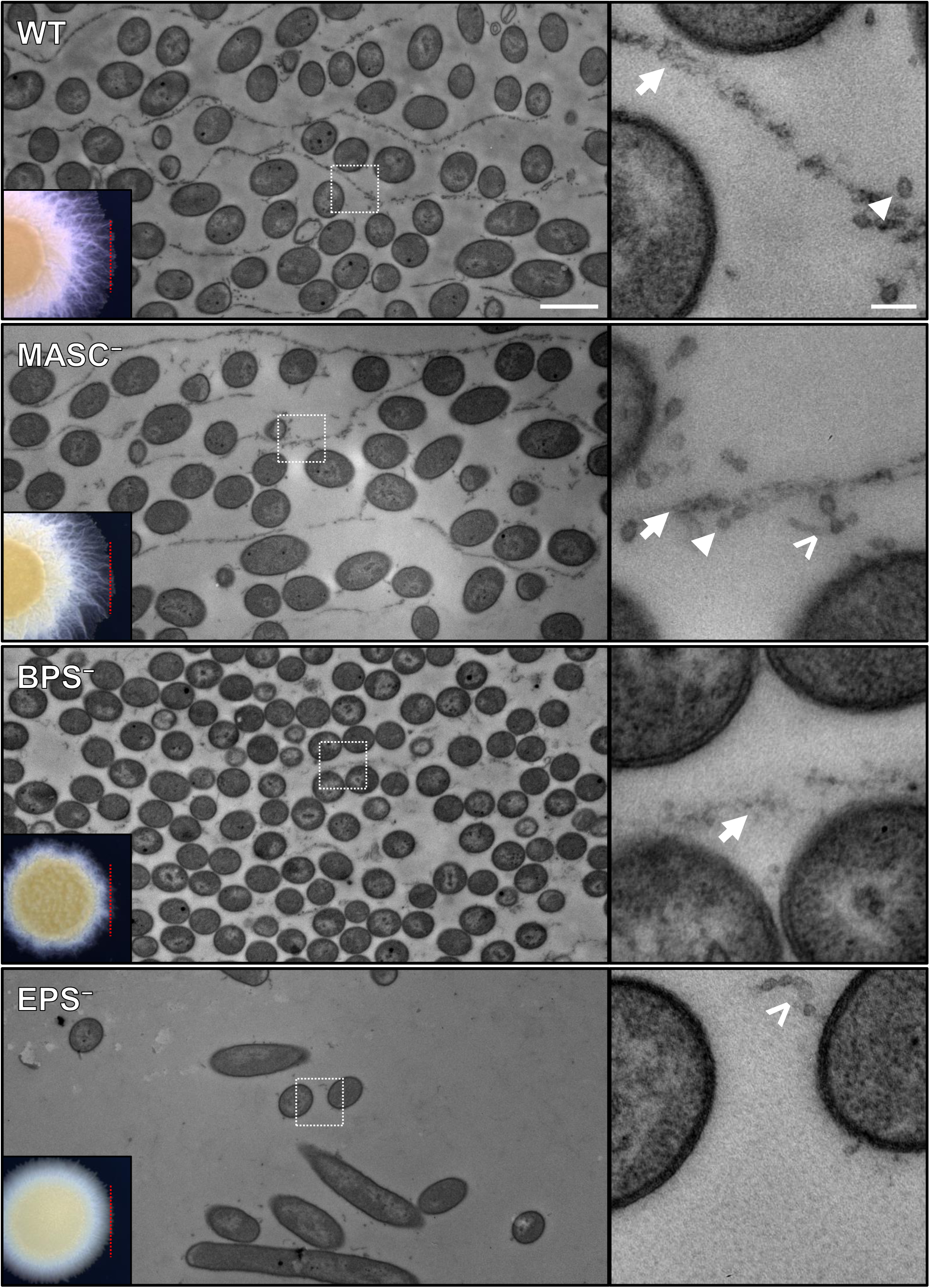
TEM analysis of Ruthenium Red-stained transverse sections cut from WT, MASC^−^ (Δ*wzaS*), BPS^−^ (Δ*wzaB*), and EPS^−^ (Δ*wzaX*) swarms. Left-side panels, *inset*: relative position of sample sectioning prior to TEM. Left-side panels: wide-angle views at 4000× magnification of internal swarm architecture. For reference, the agar substratum and the apical face of the swarm were located at the bottom and top (respectively) of each image. Scale bar: 1 µm. Right-side panels: magnified views (20 000×) of the corresponding zone within white hatched boxes in the left-side panels. Scale bar: 100 nm. Arrows (→) denote wispy putative polysaccharide-like material. Filled arrowheads (⊳) denote OMVs. Chevrons (>) denote OMV chains.

Higher-magnification views of the horizontal electron-dense ribbons revealed these structures to be of heterogeneous composition; in addition to wispy material which could represent one or more accumulated polysaccharide species, enrichments of individual OMVs as well as OMV chains were also observed at these sites (**Fig. 4**).

Taken together, these data suggest that the horizontal electron-dense structures (**Fig. 4**) are not required for nematic alignment of cells in these swarms. Furthermore, the close packing of cells in BPS^−^ swarms is consistent with BPS^−^ cells being more highly aggregative (**Fig. 2A,B**), forming fruiting bodies at lower initial cell densities (32), and displaying more compact surface EPS glycocalyces lacking fibril structures (**Fig. 2F**). Finally, the accumulation of various types of material at these horizontal electron-dense ribbons (**Fig. 4**) raises the possibility that these striations are, in essence, exclusion boundaries between different layers of *M. xanthus* cells within a swarm. Further comment on the nature of these horizontal electron-dense structures can be found in the Discussion below.

## DISCUSSION

Originally referred to as “slime fibrils/fibers”, sinewy structures connecting the surfaces of clustered *M. xanthus* cells have been known for >40 years (88, 89), with the only known requirement being the presence of the cell-surface EPS glycocalyx (55, 63, 74, 75, 78). Given the phenotypic, biochemical, and biophysical data presented herein, we propose that it is not simply the presence of cell-surface EPS that is required to mediate these inter-cell connections in *M. xanthus*, but rather the activation state of the EPS glycocalyx induced by the effects of secreted BPS.

Unfortunately, confusion exists throughout the scientific literature on the use of the abbreviation “EPS”, especially for *M. xanthus* research. Various laboratories (including ours) use “EPS” to specifically denote the principal matrix polysaccharide assembled and secreted via the WzxX-WzyX-WzcX-WzeX-WzaX proteins (32, 33, 65). However, “EPS” has also been used to non-specifically refer to diverse secreted polysaccharides (i.e. “exopolysaccharides”). For many bacteriologists, “EPS” has even more broadly come to signify “extracellular polymeric substances”, a term that has come to encompass not only secreted polysaccharides, but also polypeptides and polynucleotides. Given theses various uses, particular attention is required when interpreting and comparing findings across diverse publications.

The data presented herein provide complementary insights into the nature of T4P-dependent group motility in *M. xanthus* swarm biofilms, as they implicate the importance of BPS on several levels. Type IV pili in *M. xanthus* are known to interact with cell-surface EPS, which is how *M. xanthus* cells in rafts are proposed to move together, i.e. a T4P from a given cell is able to interact with the surface EPS layer on an adjacent cell, triggering T4P retraction, close cell–cell association, and group movement (26). Previously, single cells were found to move via T4P extension and retraction on polystyrene surfaces, but only in the presence of a viscous solution of 1% methylcellulose (90, 91). To what exactly then does a *M. xanthus* T4P bind? Though BPS^−^ cells can extend a T4P (**Fig. 2C**), these apparatus were typically shorter, thinner, and less prevalent than those in WT cells, and it is not known if pili from BPS^−^ cells are still able to interact with the “non-activated” EPS on adjacent BPS^−^ cells. Simplistically, the T4P may need to get stuck within the activated EPS matrix in WT cells, something which might not be possible in BPS^−^ cells. Alternatively, if unable to bind, this could signify that a specific motif on “activated” EPS needs to be recognized by the T4P, and that this motif is not exposed on the glycocalyx of BPS^−^ cells. Rather than the “stuck-in-goo” hypothesis, additional evidence supports the latter theory. Specifically, that (i) T4P retraction can be triggered by amine-containing polysaccharides (26), and (ii) single-cell T4P motility is possible in an aqueous microfluidic channel atop glass functionalized with a molecular coating of carboxymethylcellulose (92). The latter principal is analogous to that used for chitosan coatings in microfluidic chambers to test single-cell gliding motility on glass substrata (24, 93).

In addition, BPS secretion alters the fundamental properties of the *M. xanthus* cell surface, impacting numerous processes. Importantly, an imbalance in the EPS:BPS secretion ratio in a given cell can alter surface adhesiveness, directly influencing spatiotemporal cell–cell interaction dynamics, and by extension, swarm biofilm architecture. A greater proportion of EPS:BPS in Δ*wzaB* cells (i.e. WT levels of EPS, no BPS) results in swarms displaying a fuzzy morphology on soft agar (32) (**Fig. 1A**). Similarly, robustly increasing the production of EPS (in Δ*difG* cells) may dilute the effect of BPS, resulting in a similar fuzzy swarm morphology (**Fig. 1A**), albeit with a larger surface area (**Fig. 1B**).

However, BPS^−^ cell traits such as increased gliding speed and more frequent reversals (relative to WT) are more difficult to interpret. Given that EPS^−^ cells are not aggregative, the apparent increased “stickiness” of BPS^−^ cells (and presumed stronger association with the substratum) is not believed to lead to appreciably more efficient substratum-coupling of the Agl– Glt machinery. As the EPS glycocalyx is likely more compacted in BPS^−^ cells, and completely absent in EPS^−^ cells, we speculate that the bulk volume occupied by the BPS-activated EPS surface layer in WT cells results in suboptimal surface coupling of the gliding machinery. Conversely, overall increased vs. decreased stickiness of BPS^−^ vs. EPS^−^ cells (respectively) may indicate a role for mechanical feedback from physical interactions with the substratum influencing properties of the Agl–Glt apparatus at bFA sites and/or the Frz chemosensory system that governs polarity reversals within the cell (94), potentially affecting reversal frequency. The combined effects of increased gliding speed and potential differences in bFA stability and/or Frz system activity may also help explain the hyper-polymertropism observed for BPS^−^ cells.

The detection of internal Ruthenium Red-labelled structures near the swarm edge presented herein provides important independent support for the overall concept of leading-edge “microchannels” reported by Berleman and colleagues (17). Differences in the numbers of cells contained within layer/channel structures could be attributable to minor variations in initial swarm inoculation (volume, cell density, etc.). However, we remain skeptical of the notion that the Ruthenium Red-labelled structures are mainly composed of EPS. Ruthenium Red is a polycationic dye with a well-documented propensity for binding to a range of anionic targets (87). Originally used as a highly-effective labelling agent for pectin (a galacturonic acid-rich polysaccharide), the dye has since been shown to also bind other anionic polysaccharides, phospholipids, DNA, and proteins (87, 95-98). Though a chemical structure has yet to be determined, the composition of *M. xanthus* EPS has been studied across four publications. In total, arabinose, galactose, *N*-acetyl-galactosamine, glucose, glucosamine, *N*-acetyl-glucosamine, mannose, *N*-acetyl-mannosamine, rhamnose, and xylose sugars have been reported (32, 77, 99, 100). However, none of these reported EPS sugars carry a net-negative charge that would favour Ruthenium Red binding; this has lead us to question whether the distinct horizontal ribbons we detected within WT and MASC^−^ swarms (**Fig. 4**), as well as the walls of self-enclosed so-called “microchannel” structures at the WT swarm edge (17), are indeed composed of *M. xanthus* EPS.

Of note, extracellular DNA (eDNA) has been previously detected in *M. xanthus* biofilms and shown to bind secreted polysaccharides as well as strengthen the extracellular matrix (101). Moreover, we have herein detected the accumulation of OMVs and OMV chains at the Ruthenium Red-stained layers, material which by definition contains phospholipids as well as phosphate groups linked to the Lipid A motif of its LPS (102) (**Fig. 4**). In addition to polysaccharide, the *M. xanthus* extracellular matrix is also abundant in protein species of largely unknown functions (77, 103). Most intriguingly, BPS may be a strong candidate for the principal Ruthenium Red-labelled substance detected in the mid-swarm horizontal ribbons reported herein (**Fig. 4**) as well as the leading-edge enclosed channel structures previously reported (17). Consider that the BPS polymer is an acidic heteropolysaccharide built of repeating tetrasaccharide units; each tetrasaccharide repeat contains a proximal *N*-acetyl-D-mannosamine, followed by three distal anionic *N*-acetyl-D-mannuronic acid sugars, with the first three sugars of each repeat being randomly acetylated (32). Furthermore, pronounced Ruthenium Red-labelled horizontal striations were not detected in BPS^−^ swarms (**Fig. 4**). In addition, compositional analysis of cell-associated sugars as well as surface-active testing of culture supernatants suggest that BPS is not bound to the cell surface and is instead secreted into the extracellular milieu (32). The combined effect of a sheet of EPS glycocalyces from adjacent cells could thus be to electrostatically and/or sterically exclude secreted anionic BPS and concentrate it at EPS-free zones between cell layers. It may then be in these levels of a swarm where BPS may fully activate (destabilize) the EPS of adjoining cells, allowing for efficient T4P-mediated swarm expansion. This could also explain why no Ruthenium Red-stained structures could be detected in EPS^−^ swarms (**Fig. 4**) (17), i.e. the absence of EPS resulted in no exclusion boundaries at which BPS could accumulate. In this manner, the lumen of swarm-edge microchannels would still contain EPS material, but the surrounding Ruthenium Red-binding wall structures would contain BPS. Ultimately, BPS-dependent stratification in swarm biofilms appears to play an important role in community organization and expansion.

Our data may also shed light on findings regarding the proposed effect of the Wzb (PhpA) tyrosine phosphatase on *M. xanthus* EPS production (49). Mori and colleagues reported that a mutant lacking this Wzb tyrosine phosphatase possessed higher levels of phosphorylated BYK protein WzeS (BtkA) and (ii) the BYK domain-containing WzcB (BtkB) (49). These proteins are now known to be a part of the MASC and BPS assembly pathways, respectively (32, 33, 47). Wzb-deficient cells also exhibited faster auto-aggregation in cuvettes compared to WT cells, and were able to aggregate earlier in development resulting in faster fruiting body formation (49). Finally, the amount of Trypan Blue dye bound by Wzb-deficient cells was 134% that of WT cells (49). Accounting for these and other data, the authors concluded that PhpA may have a negative regulatory effect on EPS biosynthesis (49). However, based on (i) the demonstrated dephosphorylation of BPS-pathway WzcB by this Wzb tyrosine phosphatase (49), the faster auto-aggregation of BPS^−^ cells (**Fig. 2A,B**), (iii) the more efficient formation of fruiting bodies by BPS^−^ swarms (32), and (iv) the marginally higher amount of Trypan Blue bound by BPS^−^ cells relative to WT (**Fig. 2D**) (32), the findings for Wzb-deficient vegetative cells are more in line with a deficiency in BPS production rather than an increase in EPS production. With respect to direct effects of Wzb on the EPS biosynthesis pathway, it remains to be seen whether the Wzb tyrosine phosphatase (which already acts on WzcB and WzeS) also acts on the recently-identified WzeX BYK shown to be essential for EPS biosynthesis (32). To date, the manner by which the Dif chemosensory pathway regulates EPS production is unknown (18). Nonetheless, regulation of the putative phosphorylation state of WzeX is an attractive target for understanding changes in EPS levels during the *M. xanthus* lifecycle.

## MATERIALS AND METHODS

### Bacterial Cell Culture

The *M. xanthus* strains used in this study are listed in **Table 1**. They were grown and maintained at 32 °C on Casitone-yeast extract (CYE) agar plates or in CYE liquid medium at 32 °C on a rotary shaker at 220 rpm. The *Escherichia coli* strains used for plasmid construction were grown and maintained at 37 °C on LB agar plates or in LB liquid medium. Plates contained 1.5% agar (BD Difco).

**TABLE 1.**

| <u>Strain Code</u> | <u>Strain</u> | <u>Genotype/ Description</u> | <u>Source or Reference</u> |
| --- | --- | --- | --- |
| TM108 | *Myxococcus xanthus* DZ2 | Wild type | Laboratory collection |
| TM469 | $\Delta wzaX$ | $\Delta mxan\_7417/epsY$ | [1] |
| TM484 | $\Delta wzaS$ | $\Delta mxan\_3225/exoA/fdgA$ | [1] |
| TM529 | $\Delta wzaB$ | $\Delta mxan\_1915$ | [1] |
| EM450 | $\Delta difE$ | $\Delta mxan\_6692$ | [2] |
| EM451 | $\Delta difG$ | $\Delta mxan\_6691$ | Laboratory collection |
| TM293 | $\Omega pilA$ | $\Omega mxan\_5783$ (Tet<sup>R</sup> cassette insertion) | Laboratory collection |

[1] Ducret A, Valignat M-P, Mouhamar F, Mignot T, Theodoly O. Wet-surface-enhanced ellipsometric contrast microscopy identifies slime as a major adhesion factor during bacterial surface motility. Proc. Natl. Acad. Sci. U. S. A. 2012. 109(25):10036-10041.
[2] Moine A, Agrebi R, Espinosa L, Kirby JR, Zusman DR, Mignot T, Mauriello EMF. Functional organization of a multimodular bacterial chemosensory apparatus. PLOS Genet. 2014. 10(3):e1004164

### Phenotypic Analysis

Exponentially-growing cells were harvested and resuspended in TPM buffer (10 mM Tris-HCl, pH 7.6, 8 mM MgSO_4_ and 1 mM KH_2_PO_4_) at the final concentration of OD_600_ 5.0 for gliding, T4P-dependent expansion, and developmental assays. This cell suspension (5 μL) was spotted onto CYE 1.5% agar, CYE 0.5% agar, or CF 1.5% agar for gliding flare, T4P-dependent swarm expansion, or developmental (i.e. fruiting body formation) analysis, respectively. Plates were incubated at 32 °C for 30 h for gliding flares, 72 h for T4P-dependent swarm expansion, and 75 h for fruiting body formation, then photographed with an Olympus SZX16 stereoscope with UC90 4K camera. Gliding flares were imaged using the 2× objective at 8× zoom, using linear colour, with illumination control wheel set halfway between the brightfield cartridge and the open slot on the illumination wheel. For T4P-dependent motility, swarms were imaged using the 0.5× objective at 1× zoom, using linear colour and darkfield illumination. For fruiting bodies, structures were imaged using the 0.5× objective at 2× zoom, using high-quality colour and oblique illumination.

### Transmission Electron Microscopy

For T4P visualization, a 50 µL drop of overnight liquid culture was transferred to a copper grid and incubated at ambient temperature for 5 min. Grids were then dried with bibulous paper, stained with 3% PTA (pH 6.0) (Mecalab) for 2 s, and dried again with bibulous paper. For swarm biofilm cross-sections, swarm samples were fixed in 2.5% glutaraldehyde in cacodylate buffer at pH 7.4 with 0.2M sucrose overnight and then washed three times with cacodylate buffer. Then post-fixed in 1.33% osmium tetroxide in Collidine buffer (pH 7.4) for 1 h and stained with 5 mM Ruthenium Red for 1 h at ambient temperature. After dehydration by successive passages through 25, 50, 75, 95% and 100% (twice) solutions of ethanol in water (for 30 min each), samples were immersed for 16–18 h in Spurr:acetone (1:1 v/v). Samples were then embedded in Spurr resin (TedPella) before incubation at 60–65 °C for 20–30 h. After polymerization, samples were sectioned (90 – 150 nm) using an ultramicrotome (LKB Brooma - 2128 Ultratome). Sections were collected on formvar / carbon-coated copper 200-mesh grids. Samples were stained with 5% uranyl acetate in 50% ethanol for 15 minutes followed by lead citrate for 5 minutes. Imaging for T4P and swarm biofilm cross-sections was carried out using a Hitachi H-7100 transmission electron microscope with AMT XR-111 camera.

### Scanning Electron Microscopy

Glass coverslip discs were first washed in 100% EtOH for 1h and left to dry at RT under sterile conditions. The cleaned discs were immersed for 1h at RT in 0.01% poly-L-lysine solution and allowed to dry. Discs were placed one per well in a 24-well polystyrene cell-repellent plate and overlaid with 2 mL of overnight CYE bacterial cultures. Plates were then covered, sealed with Parafilm, and incubated overnight with shaking at 32 °C. After incubation, the media was removed and the cells attached to the coverslips were fixed in 1 mL of 2.5% glutaraldehyde in 0.1 M cacodylate buffer (pH 7.4) for at least 1 h and washed three times in 1 mL of 0.2 M cacodylate buffer for 5 min. After post-fixation in 500 µL of 1.33% osmium tetroxide (in 0.2 M cacodylate buffer) for 1 h, bacteria were dehydrated with increasing ethanol concentrations (25, 50, 75, 95 and 100%). From the 100% EtOH bath, coverslips were critical-point-dried using CO_2_ (Leica EM ACE600), coated with 3 nm gold/palladium (Leica CPD300) and examined with a JEOL JSM-7400F scanning electron microscope (3 kV-LEI detector).

### Single-Cell Gliding Motility Analysis

For phase-contrast microscopy on agar pads, cells from exponentially-growing cultures were sedimented and resuspended in TPM buffer (OD_600_ 0.7), spotted (3 µL) on a glass coverslip, and overlaid with a 1.5% agar pad prepared with TPM buffer. For motility analysis, cells were left to adhere for 5 min prior to imaging at 32 °C. Images were obtained using an Axio Observer 7 microscope, with a Plan Apochromat 40× 1.3 oil objective VIS-IR M27 (for total magnification 400×), an Axiocam 512 as camera, and a TL LED as light source (Zeiss). Images were taken at 30 s intervals. The microscope was operated using the Zen 2.6 Pro software suite (Zeiss).

Prior to analysis, image stacks were treated in FIJI as follows to optimize tracking: *Step 1:* Enhance Contrast (0.3%), Normalize, Process All Slices. *Step 2:* Subtract Background (Rolling ball radius of 15.0 pixels, Light background, Process all slices). *Step 3:* Image alignment in stack via StackReg (Translation) plug-in. Cell gliding speeds were then calculated using the MicrobeJ module for FIJI (104): *Step 1:* Under the Bacteria tab, Tracking Parameters were adjusted (Max Entropy, Medial Axis, Area: 18-800, Length: 10-max, Width: 1.5-max), with only cells tracked for a minimum lifespan of 20 frames used for analysis. *Step 2:* Automatically-detected objects were manually curated to remove instances of object merging, background artifact detection, and reference cell switching, followed by comparison of Velocity Means (pixels/frame). *Step 3:* Gliding speeds were converted to “µm/min”. Reversals of gliding direction for these tracked cells were manually counted, with a minimum displacement of ∼75% of cell length considered a reversal.

### Polymertropism Testing

Aspect ratio (AR) vs. time analyses were modified from a published report (84) and were performed as previously described (85). Cells of *M. xanthus* (grown in CYE at 28 °C to ∼ 5 × 10^8^ cells/mL) were sedimented (4000 × *g*, 10 min), then resuspended in CYE broth to 5 × 10^9^ cells/mL, and used to inoculate (4 µL) compressed and uncompressed round 85 mm CTTYE agar plates. An ∼ 1 cm length of 5.56 mm outer-diameter Tygon tubing was inserted against the plate wall to compress the agar (84), with cells on these plates inoculated 43 mm from the inserted tubing. Following incubation at 30 °C for 24, 52, 90, 120, and 144 h, colony perimeters were marked at each interval. The AR of each swarm was then calculated for each time point by taking the quotient of the colony width and colony height; a round swarm will produce an AR near-or-equal to one, whereas an elongated swarm will produce an AR > 1. For each replicate dataset, linear best-fit lines were plotted, followed by determination of the slope (i.e. AR/time). Average slope values were calculated for each strain and normalized as a percentage of the AR/time for the WT strain.

### Auto-Aggregation Testing

Using a modified version of an auto-aggregation protocol (60), overnight *M. xanthus* cultures (10 mL) were sedimented in 15 mL conical tubes (4000 × *g*, 5 min), followed by resuspension of pellets in TPM buffer (10 mL) and OD_600_ determination using disposable cuvettes. Specific resuspension volumes were aspirated and sedimented in a microfuge tube (4000 × *g*, 5 min); pellets were resuspended in 1 mL CYE broth or TPM buffer to a final OD_600_ of 0.5, followed by transfer to a polystyrene spectrophotometer cuvette. Samples were vigorously aspirated/ejected in the cuvette for 10 s using a p200 micropipette, followed by immediate reading of the OD_600_ (*t* = 0). Subsequent OD_600_ readings were obtained at 10 min intervals up to 150 min of monitoring. Finally, all OD_600_ readings were normalized to the OD_600_ determined at *t* = 0 for each sample.

### Cell-Surface Hydrophobicity Testing

To analyze relative differences in cell-surface hydrophobicity, we employed a modified version of the classic microbial adhesion to hydrocarbons (MATH) assay (71, 72). Based on the OD_600_ of *M. xanthus* overnight cultures (12.5 mL CYE) measured via NanoDrop 2000c spectrophotometer (Thermo), sufficient culture volume was removed, sedimented (6000 × *g*, 5 min) in 2 mL conical tubes, followed by pellet resuspension in 4 mL fresh CYE medium via using a p1000 micropipette to a final OD_600_ of 1.0. The OD_600_ of the equilibrated 4 mL resuspensions (i.e. Mix 1) was read in a quartz cuvette. Cell suspensions were then transferred to a new 15 mL conical tube using a p1000 micropipette, mixed via vortex (maximum speed, 20 s), then transferred back to the quartz cuvette for OD_600_ determination (i.e. Mix 2); this step was performed as an internal control to ensure that downstream changes in OD_600_ were not simply due to further mixing of the sample, particularly for hyper-aggregative strains. Samples were returned via aspiration with a p1000 micropipette to the same 15 mL conical tube, followed by addition of 300 µL hexadecane (Sigma). To generate emulsions, cell–hydrocarbon mixtures were blended via vortex (maximum speed, 20 s), then rapidly transferred back to the quartz cuvette using a p1000 micropipette (i.e. Mix 3). The OD_600_ of this resuspension was immediately determined (*t* = 0), followed by readings at 5 min intervals for the next three data points. After the initial 15 min of monitoring, emulsion separation was further monitored at 10 min intervals to a final monitoring time of 65 min. All OD_600_ readings were normalized to the initial OD_600_ determined for samples at the “Mix 1” stage of processing.

### Trypan Blue Dye Retention

Trypan Blue dye-retention analysis was performed as previously described (32). In brief, cells grown overnight in CYE cultures were resuspended to OD_600_ 1.0 in TPM. Resuspended cells or a cell-free blank (900 µL) were added together with Trypan Blue stock solution (100 µL) to a microfuge tube, then briefly pulsed (1 s) on a vortex mixer. Samples were incubated at room temperature, in an aluminum foil-covered tube rack, on a rocker platform (1 h) to permit dye binding by the cells. Samples were then sedimented (16 000 × *g*, 5 min), followed by transfer of the top 900 µL of blank or clarified supernatant to a disposable spectrophotometer cuvette. Using the cell-free “TPM + Trypan Blue” sample, the spectrophotometer was blanked (585 nm). For each clarified supernatant, the absorbance at the same wavelength (A_585_) was determined. Absorbance values were then normalized to A_585_ for the WT sample.

## ACKNOWLEDGEMENTS

The authors would like to thank (i) Éric Déziel for insightful discussions and troubleshooting regarding hydrophobicity testing and (ii) Philippe Constant for valuable input on biostatistics. A Discovery operating grant (RGPIN-2016-06637) from the Natural Sciences and Engineering Research Council of Canada and a Discovery Award (2018-1400) from the Banting Research Foundation fund work in the lab of S.T.I. as well as studentships for F.S. and N.Y.J.; both are also recipients of graduate studentships from the PROTEO research network. The funders had no role in study design, data collection and interpretation, or the decision to submit the work for publication.

## AUTHOR CONTRIBUTIONS

FS and STI conceived of and planned the study.

FS and NYJ performed stereoscopic phenotypic analyses and measured colony surface areas.

AN, FS, and NYJ prepared samples for electron microscopy

AN, FS, and NYJ performed transmission electron microscopy, while the former two performed scanning electron microscopy.

FS performed dye-binding assays.

STI performed auto-aggregation analyses. FS tested strain hydrophobicity.

NYJ quantified cell motility and cell reversals.

DJL tested polymertropism responses, with analysis by STI and DJL.

FS, NYJ, and STI wrote the manuscript.

FS, NYJ, and STI generated figures.

STI, AGG, and FJV contributed personnel and funding support.

